# Clonotypic Heterogeneity In Cutaneous T-Cell Lymphoma Revealed By Comprehensive Whole Exome/Transcriptome Sequencing

**DOI:** 10.1101/405415

**Authors:** Aishwarya Iyer, Jordan Patterson, Thomas Salopek, Gane Ka-Shu Wong, Robert Gniadecki

## Abstract

Mycosis fungoides (MF), the most common type of cutaneous T-cell lymphoma, is believed to represent a clonal expansion of a transformed skin resident memory T-cell. T-cell receptor (TCR) clonality (i.e. identical sequences of rearranged TCRα, β and γ), the key premise of this hypothesis, has been difficult to document conclusively because malignant cells are not readily distinguishable from the tumor infiltrating, reactive lymphocytes, which contribute to the TCR clonotypic repertoire of MF. Here we have successfully adopted the technique of targeted whole exome and whole transcriptome sequencing (WES/WTS) to identify the repertoire of rearranged TCR genes in tumor enriched samples from patients with MF. Although most of the investigated biopsies of MF had the expected monoclonal rearrangements of TCRγ of the frequency corresponding to the frequency of tumor cells, in half of the samples we detected multiple (up to seven) TCRα and -β clonotypes by WES and WTS. Our findings are compatible with the model in which the initial malignant transformation in MF does not occur in mature, memory T-cells but rather at the level of T-lymphocyte progenitor after TCRγ rearrangement but before TCRβ or TCRα rearrangements. The WES/WTS method is potentially applicable to other types of T-cell lymphomas and enables comprehensive characterization of the TCR repertoire and mutational landscape in these malignancies.

## Introduction

Mycosis fungoides (MF) is the most prevalent form of cutaneous T-cell lymphoma (CTCL). In early stages it presents with scaly plaques on the skin which may progress into tumors and finally disseminate to lymph nodes and to other organs.^1–3^ MF can be viewed as a model of low-grade T-cell lymphomas: it has a chronic, relapsing course, low-grade proliferation, chemotherapy resistance and 5-year mortality approaching 50%.^1,4^ MF expresses markers of memory T-cell and appears to exhibit T-cell receptor (TCR) monoclonality and is thus considered to be caused by malignant transformation of a mature T-cell residing in the skin.^5^

TCR is an excellent marker of T-cell lineage because TCR-δ, -γ, -β and -α loci become sequentially rearranged during intrathymic maturation of T-cell from diverse V, (D) and J gene segment pools, and the unique products of the rearrangements are retained (with a notable exception of TCR-δ) in all daughter cells.^6^ Complementarity-determining region 3 (CDR3) encoded by the V(D)J junction is especially useful for lineage tracing because its sequence heterogeneity is increased beyond the combinatorial V(D)J diversity by random insertions and deletions of nucleotides during segment recombination.^7^ Thus, identical TCRγ, -β and -α sequences of CDR3 in all lymphoma cells would be a conclusive proof that malignant transformation took place in a mature T-cell which had completed TCR rearrangement. However, true TCR monoclonality, as defined by a single T-cell clonotype, has not been demonstrated in CTCL. Usually, the dominant clone is accompanied by several other TCR clones thought to originate from reactive, tumor-infiltrating T-cells. Statistical methods have been used to formally determine clonality^8^ but these methods neither distinguish between tumor clones and expanded reactive clones nor determine clonotypic heterogeneity of the tumor itself.

Determination of the clonotypic structure of CTCL is practically important, because clonality assessments are used for clinical diagnosis, prognosis and staging of CTCL.^1,9^ The most widely used method based on multiplexed PCR amplification of TCRγ and -β and Genescan analysis^10^ is currently being replaced by methods based on high-throughput sequencing of PCR amplified CDR3 regions.^9,11–13^ They seem to have superior sensitivity and specificity in the detection of the tumor clone but they cannot differentiate CDR3 sequences derived from tumor cells versus those derived from reactive T-cells and do not provide any measure of sample purity (the percentage of neoplastic cells). Moreover, the amplification step with multiplex PCR makes sequencing of the complex TCRα locus virtually impossible. Sequencing of TCRα can be achieved by RNA-seq where primers binding to the invariable constant TCR segment are used but only the transcribed TCR alleles are detected and the information on other non-productive rearrangements in the genome is not captured. Unfortunately, RNA-seq results may be distorted by presence of alternatively spliced mRNA and allele silencing, not uncommonly seen in cancer.^11^

It has been reported that the CDR3 sequences of rearranged TCRβ genes can be retrieved from the whole exome sequencing (WES).^14^ Based on this finding we have developed a protocol in which samples are analysed by the probe capture WES and the whole transcriptome sequencing (WTS). This allowed us to identify recombined TCRα, -β and -γ sequences from DNA in MF patients and compare their respective expression patterns. Since WES also allows to quantify the percentage of tumor cells in the sample, we were able to reconstruct clonotypic composition of MF and provided evidence for TCR heterogeneity of this lymphoma.

## Results

### Identification of T-cell clonotypes from whole exome sequencing (WES) and whole transcriptome sequencing (WTS)

To determine whether sequences of CDR3 regions and TCR clonotypes can be determined from WES and WTS we performed laser capture microdissection of the areas of atypical lymphocytic infiltrate in 14 biopsies of plaques (early lesions) and tumors (advanced lesions) of 9 patients with MF (**Fig 1**). To be able to compare the results of WES and WTS directly, we purified DNA and RNA simultaneously from the same samples. As shown in **Fig 2 A, B** (and in supplementary **Table S1**), the capture based WES technique successfully identified numerous CDR3 sequences corresponding to TCRα, TCRβ and TCRγ clonotypes. Using the sequencing depth of 87×10^6^ reads we were able to capture (median and range): 143.5 (66-393) TCRα, 40 (13-95) TCRβ and 6 (2-15) TCRγ clonotypes. The relative excess in TCRα abundance is readily explainable by the fact that during T-cell development the TCRβ is under the strict allelic exclusion, but TCRα locus is usually rearranged on both chromosomes, sometimes in multiple rounds resulting in 2-4 TCRα rearrangements per single TCRβ rearrangement.^15^ This explanation is confirmed by WTS results documenting a comparable number of expressed TCRβ clonotypes to the number of clonotypes identified at the DNA level (35.5 vs 40) and practically the same median number of transcribed TCRα clonotypes (n=50) **(Fig 2 B)**. There was no bias in V and J segment detection in the control samples of peripheral blood (supplementary **Fig S1**) with our WES protocol.

**Fig 1.**
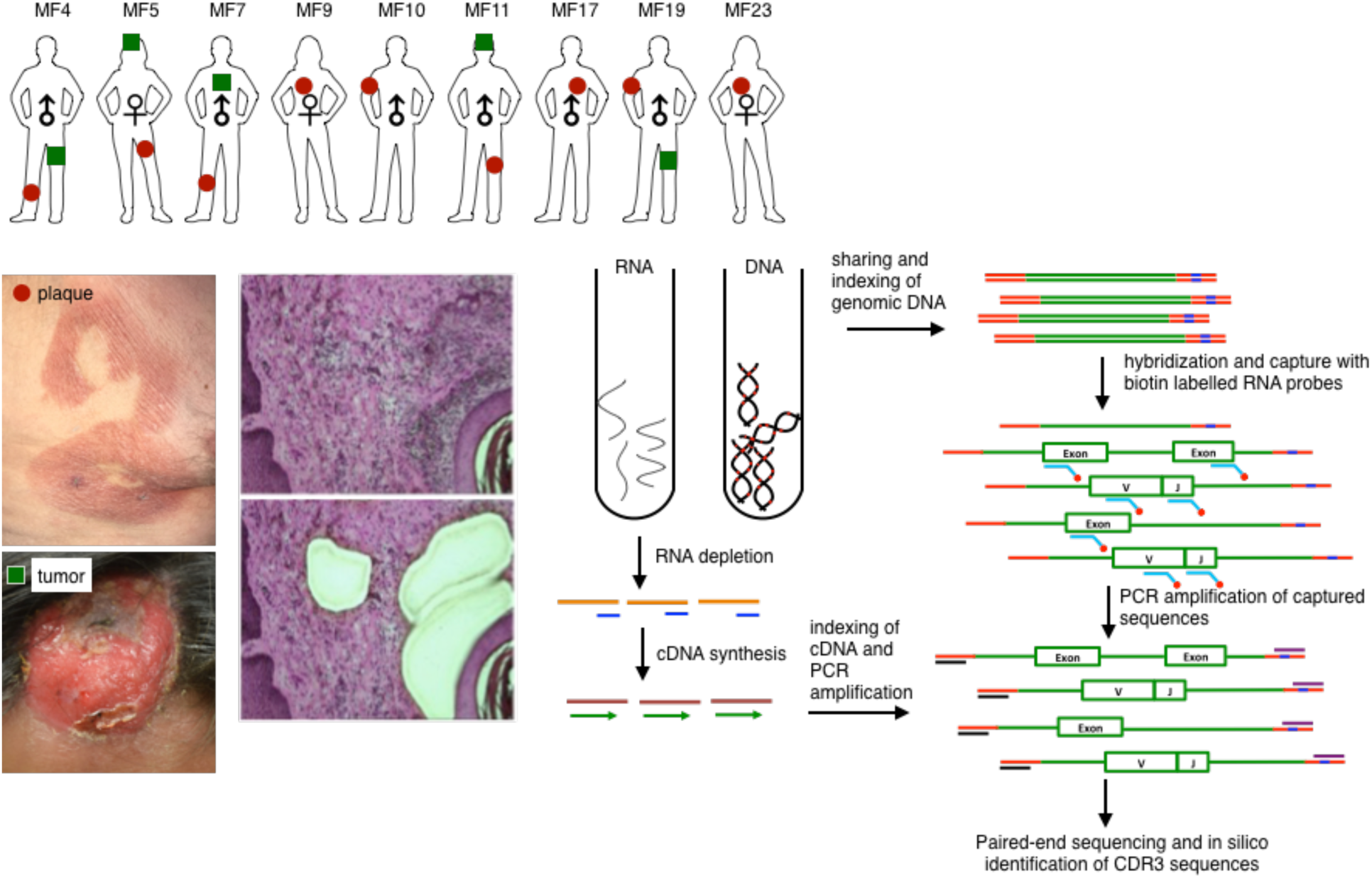
Schematic representation of sample collection, processing and TCR sequencing. 4mm punch biopsies were collected from 9 patients with MF, from early lesions (plaques, red circle) or tumors (green squares). Biopsies were cryosectioned and laser microdissected to capture tumor cells. DNA and RNA was isolated simultaneously from the microdissected material and processed for whole exome sequencing (WES) and whole transcriptome sequencing (WTS). WTS data is unavailable for samples MF5P, MF9, MF10, MF17, MF23. Not shown are control samples: isolated CD4+ T-cells from 4 healthy donors (WTS) and whole blood from patients MF5 and MF11 (WES). Indicated in green is the gene sequence, red is the adapter sequence and blue is index sequence.

**Fig 2.**
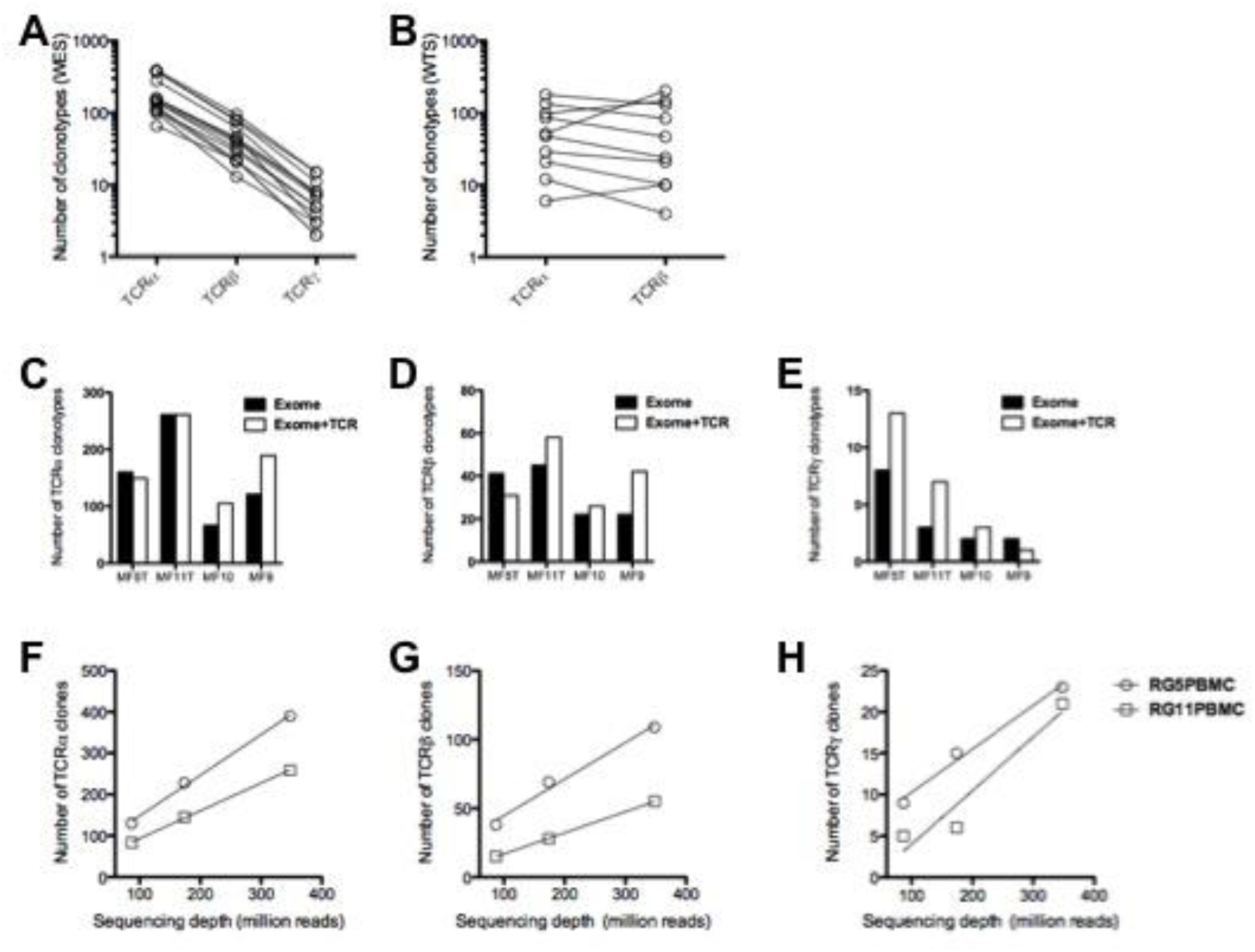
Efficiency of the WES probe capture and WTS protocols in detection of CDR3 clonotypes in biopsies from MF. All The samples were sequenced using whole exome probe capture **(A)** and WTS **(B)** and the number of clonotypes was determined for each sample for TCRα, TCRβ and TCRγ, as indicated. The lines connect the results for the same sample. **C-E:** The effect of TCR-specific probes. The capture was performed in four samples with whole exome probes as in A (Exome), or with whole exome probes combined with specific TCR capture probes (Exome+TCR), sequenced and the number of unique clonotypes for TCRα (C), TCRβ (D) and TCRγ (D) was determined, as in panel **A**. The addition of probes slightly increased the number of TCRγ clonotypes (p=0.024, paired t-test), but not of TCRα or TCRβ. **F-H:** The effect of sequencing depth on clonotype detection. Two samples of whole peripheral blood mononuclear cells were sequenced with WES at a maximum of 400 million reads as in **A**. The saturating sequencing depth in identifying TCR sequences was 348 million reads.

### Efficiency of probe capture technique in identification of T-cell clonotypes

Previous protocols using gene capture and high throughput sequencing used TCR specific capture probes rather than the vast panel of probes for the entire exome.^16^ The drawback of that approach is that use of fewer probes can lead to decreased capture selectivity. Since the exome capture probe set was not specifically designed to capture TCR genes, we asked whether efficiency of capture can be increased by additional probes targeting V and J segments of TCRα, β and γ. As shown in **Fig 2 C-E,** those additional probes increased the total number of identified clonotypes in 3 of the 4 samples tested, but the difference was not statistically significant. Therefore, for the subsequent experiments we used standard exome capture probes. We have also tested the sequencing depth on clonotype detection efficiency by sequencing two total blood samples with 400 million reads each. It was observed that the current capture protocol reached a saturation in identifying unique T-cell clonotypes at 348 million reads. The efficiency with increased sequencing depth still remained highest for TCRα and lowest for TCRγ (**Fig 2 F-H**).

### Analysis of malignant TCR clonotypes in MF

MF is thought to develop from memory T-cells and therefore it should have the same TCRγ, -β and -α clonotype. The concept of monoclonality of T-cell lymphoma has been well documented using multiplex/heteroduplex PCR amplification and detection by capillary electrophoresis or high throughput sequencing^9,12^ and is used as a diagnostic test in CTCL. We were therefore interested whether our method of WES/WTS clonotype detection can identify those TCR clones in MF samples. The biopsies always contain variable, usually unknown, amounts of reactive T-cells contributing to the repertoire of TCR clonotypes, but the clonotypes of frequency exceeding 15% are usually confidently classified as monoclonal.^9,12^ As shown in **Fig 3**, if the 15% clonotype frequency threshold is applied, only 6 of 14 MF can be classified as monoclonal on the basis of TCRβ clonotypes identified from WES. The frequency of the most abundant TCRβ clonotype was higher when WTS was used (6 of 9 samples). In WES analysis of TCRα only 3 of 14 samples could be classified as monoclonal, but 6 of 9 biopsies showed >15% of the most frequent clonotype with WTS.

**Fig 3.**
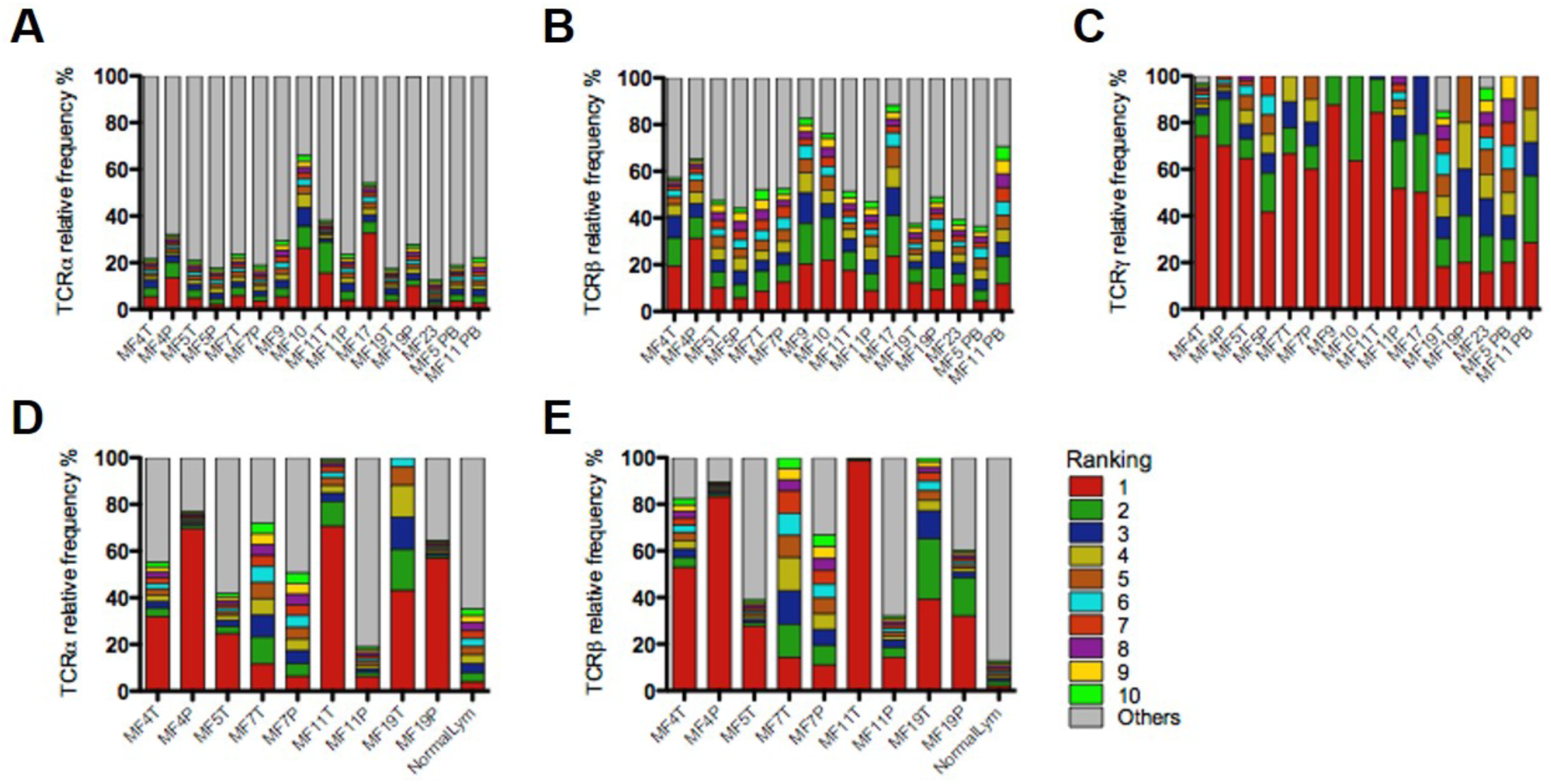
Relative frequency of T-cell clonotypes. A-C: TCRα (**A**), TCRβ **(B)** and TCRγ **(C)** repertoire sequences identified from WES of MF samples. Sample ID relates to patient number as in Fig 1 with the suffix P (plaque), T (tumor) or PB (peripheral blood mononuclear cells). **D-E:** TCRα (D) and TCRβ (E) repertoires identified by WTS of MF samples. NormalLym: pooled CD4+ normal lymphocytes from 4 healthy donors. Each bar represents an individual CDR3 amino acid clonotype, with red and moss green indicating the first and tenth ranked clonotype in a decreasing order of relative frequency. Grey bars represent the the rest of the identified clonotypes in the samples.

The WES information can also be used to determine the copy number aberrations in cancer genome and hence calculation of the enrichment tumor cells in the sample. We have intentionally decided to used microdissected samples because it is well known that the ratio of malignant cells to reactive T-cells in the lesional skin in MF may be low, especially in the early plaque lesions. Even in the microdissected samples the proportion of malignant cells varied between 43.1% to 88.1% (median 58.5%) and there were no differences between the plaques and the tumors. Contrary to expectation, neither the frequency of the most abundant (dominant) clone nor diversity index (inverse Simpson index) were correlated with the proportion of tumor cells in the sample (with an exception of a weak correlation between TCRα and tumor DNA, R^2^=0.50, p=0.021), **Fig 4A** (and **supplementary Figure S2**). More surprising was the finding that a single TCRβ clonotype cannot account for all malignant cells in the sample **(Fig 4B)**. Even in samples where the ratio of the dominant TCRγ clonotype to the proportion of tumor cells ∼1 (MF4T, MF5P, MF5T, MF7T, MF7P, MF11P, MF11T), i.e. samples with perfect TCRγ monoclonality, the dominant TCRβ clonotype could account for only 20.7% of tumor cells. As can be seen in **Fig 3B**, WES revealed presence of additional one to three TCRβ clonotypes which had a comparable frequency to the dominant clonotype. Intriguely, WTS for these samples revealed single dominant TCRβ and TCRα in MF4T, MF5T and MF11T, oligoclonality in MF7T, MF7P and policlonality for MF11P (**Fig 3**). This result illustrates that a malignant T-cell clone defined by identical TCRγ can rearrange multiple TCRβ and in some instances also express more than a single TCRα and -β mRNA.

**Fig 4.**
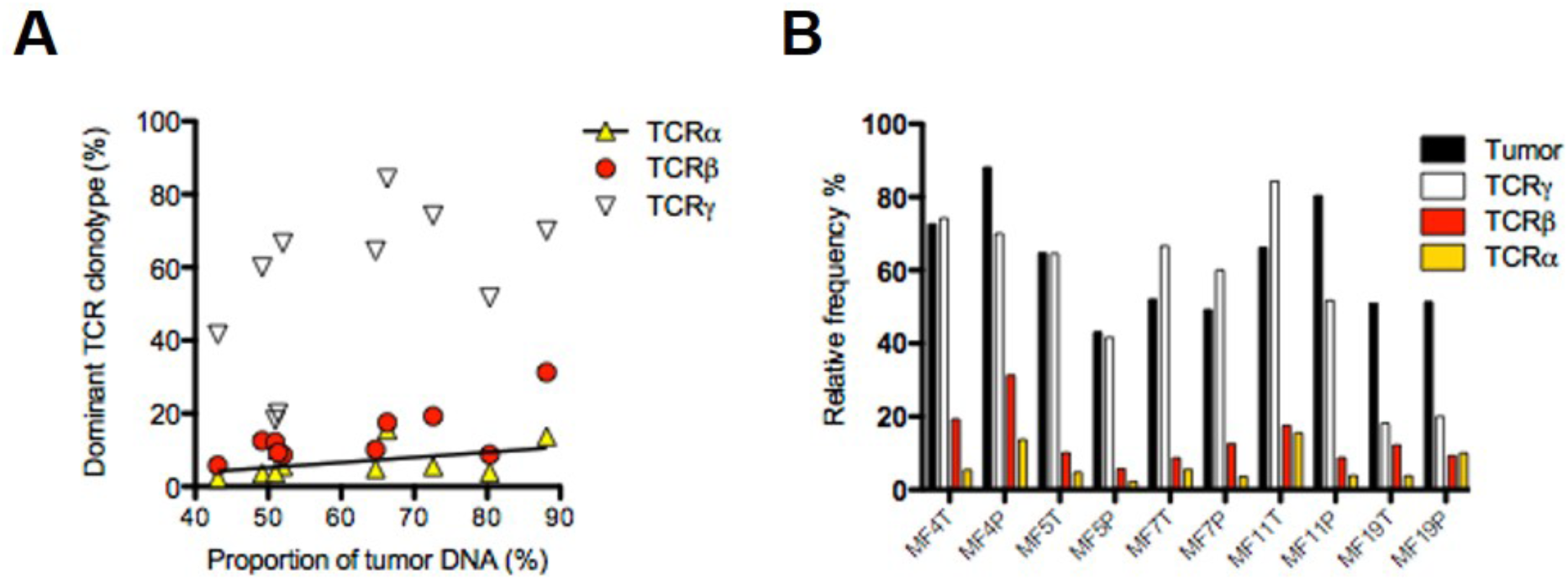
Clonotypic diversity of MF. **A**: In each sample the percentage of tumor DNA was determined from WES data and plotted against the frequency of the dominant TCRα, -β or -γ clonotype (i.e. the clonotype that ranked 1 as in Fig 3). There was no correlation between those values for TCRβ or -γ and a weak correlation for TCRα (R^2^=0.50, p=0.021 - regression like is shown). **B:** Contribution of the dominant clonotypes of TCRα, -β or -γ relative to the tumor DNA enrichment of the sample. Note that for samples MF4T, MF4P, MF5T, MF5P, MF7T, MF7P and MF11T the proportion of the dominant TCRγ clonotype equals approximately the proportion of tumor DNA in the samples indicating that all tumor cells share the same TCRγ clonotype. However, in the same samples the relative proportion of the dominant TCRα and -β clonotypes are only 21.5% (range 13%-44.6%) indicating that other clonotypes are found in tumor-derived DNA.

### Identification of shared TCR clonotypes

Simultaneous exome and transcriptome sequencing allowed us to compare both techniques for clonotype detection in MF. With the exception of MF5T and MF7P, maximum of 2-5 TCR clonotypes were found to be shared for WTS and WES for a given sample (**Fig 5A**). Regardless the cause of the lower than expected degree of overlap between WES and WTS (e.g. due to mutations, non-productive DNA rearrangements, alternative RNA splicing, incomplete intronic exclusion) this finding underscores the value of using both DNA and RNA as a source for clonotypic analysis in CTCL.

**Figure 5:**
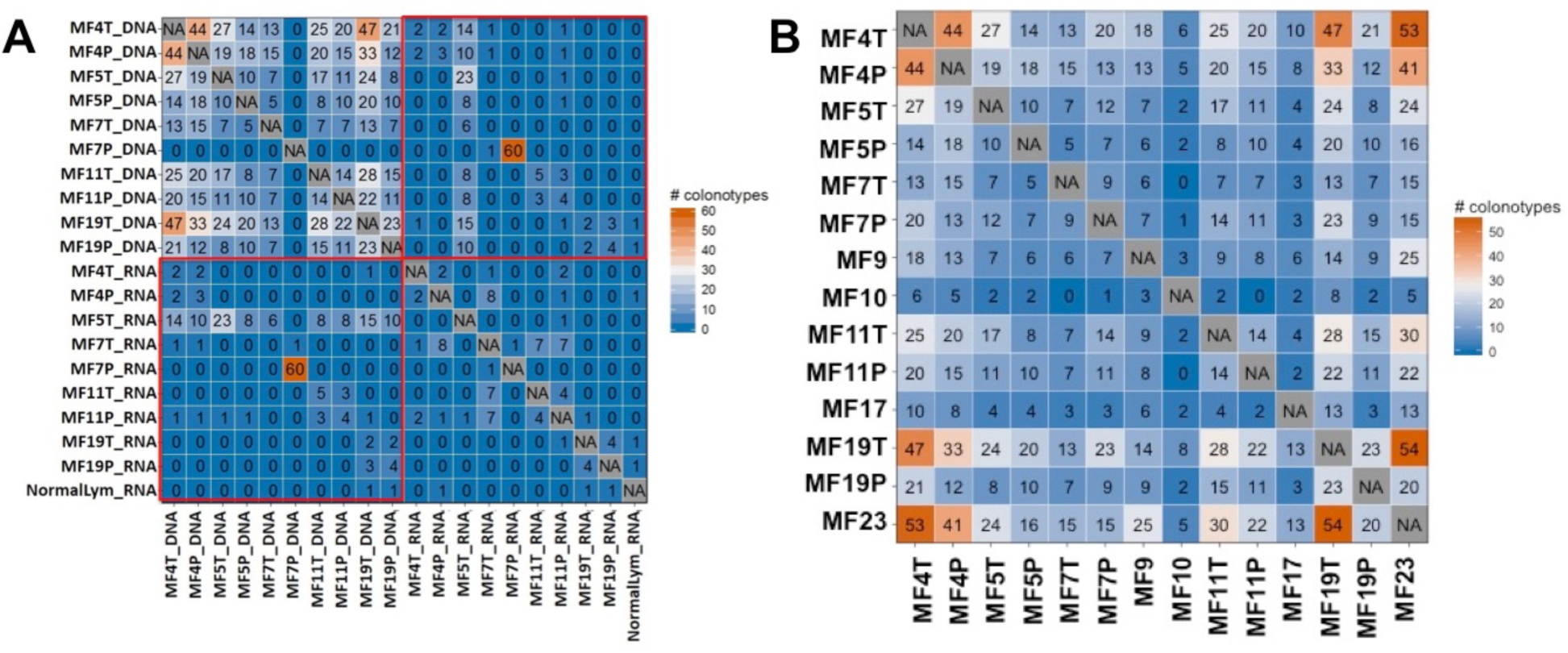
Shared T-cell clonotypes. CDR3 sequences identified using whole exome sequencing (WES) and whole transcriptome sequencing (WTS) were tested for overlap. As highlighted in the upper right or lower left quadrant of the heatmap in **A.** the highest degree of overlap was observed in MF7P. Also, the lower left quadrant indicates the shared T-cell clonotypes between the MF samples identified using WES. **B.** Indicates the common TCR sequences identified using WES. This is also represented in the upper right quadrant of panel **(A)** heatmap except for samples MF9, MF10, MF17 and MF23. The degree of overlap is indicated by the number and colour in each cell of the heatmap.

Given the vastness of the CDR3 repertoire it could be expected that individual clonotypes are not shared in samples from different patients. However, interindividual clonotype sharing was relatively common with the highest number of 54 shared clonotypes shared between MF19T and MF 23 (**Fig 5B**). The number of shared clonotypes was lower for WTS where up to 8 identical clonotypes were found in different patients **(Fig 5A** and supplementary **Table S3**). The Vα and Vβ segment usage did not reveal any clues as to the functional role of those clonotypes with two exceptions: high representation of pseudogenes (TRAV11, TRAV28, TRAV31, TRBV12-1, TRBV22-1) and the ability of TCR composed of TRAV3 (and its paralog TRAV8-4) and TRBV3-1 in recognition of lipid antigens via CD1a.^17^

## Discussion

In this report we demonstrate that TCR repertoire in CTCL can be assessed by probe capture WES, and WTS using samples such as microdissected cells. The advantage of our approach is integration of the TCR repertoire data with cancer-related mutations and the transcriptome that allows for precise calculation of the percentage of malignant T-cells in the sample. The method is especially useful for identification of TCRα locus rearrangements that do not amplify reliably with multiplex PCR due to large number of V and J genes. To date, all data on TCRα were gathered with RNA sequencing^18,19^ and very little is known about the diversity of TCRα at the DNA level.

The major drawback of our method is its lower robustness than the PCR-based methods in capturing the whole TCR repertoire in the sample. WES/WTS yielded hundreds rather than thousands TCRα and TCRβ clonotypes which although sufficient to analyse TCR rearrangement in tumor cells does not allow for comprehensive estimation of the entire T-cell diversity. The number of detected clonotypes was linearly dependent on sequencing depth but the method achieved saturation at the depth of 348 million reads where a maximum of 390 TCRα and 109 TCRβ clonotypes could be detected in human full blood. WES was least sensitive for TCRγ and did not yield the expected number of clonotypes even at the saturating coverage, which suggested that limited identification of TCRγ rearrangements was due to inefficient capture rather than the sequencing coverage. It is likely that decreasing the number of exome capture probes will reduce probe competition and increase the number of identifiable clonotypes.

Analysis of TCR repertoire in MF by WES/WTS led to unexpected conclusions regarding the nature of clonal expansion of malignant cells. By comparing the proportion of tumor-derived DNA in the sample with the relative frequencies of TCRβ and TCRα clonotypes we found evidence for existence of multiple, rather than single, malignant T-cell clonotypes. Especially informative were the cases where the proportion of monoclonal TCRγ rearrangement matched the proportion of tumor-derived DNA, indicating that the sample was composed of a population of malignant cells sharing identical TCRγ clonotype (e.g. cases MF4T, MF4P, MF5P, MF7T, MF7P, MF11T, **Fig 3, 4**). Instead of expected TCRβ monoclonality, we detected 2-7 TCRβ clonotypes and multiple TCRα clonotypes in WES. In MF7T and MF7P also contained multiple expressed TCRβ clonotypes. This indicates that at least in some cases of MF, the initial transformation does not happen at the level of skin-resident memory T-cell but much earlier during lymphocyte development, i.e. after completion of TCRγ rearrangement but before initiation of TCRβ and -α recombination. Thus, all malignant cells inherit the identical TCRγ CDR3 sequences, but not TCRβ or TCRα which would be different in the subclones descending from the same precursor. Other groups that performed TCR sequencing in CTCL also found evidence of oligoclonality ^20^. Recently, Ruggiero et al.^18^ using ligation-anchored PCR mRNA amplification and sequencing of TCRα and TCRβ in Sézary syndrome found oligoclonal, rather than monoclonal pattern in 4/10 patients and polyclonal TCR repertoire was reported in subgroups of patients with PTCL-NOS or AITL^19^. Supportive evidence comes also from the studies showing multiple TCRβ transcripts in CTCL with the copy number aberration of chromosome 7 containing TCRβ.^21^ Because recombination activating genes RAG-1 and RAG-2 are not active in mature T-cells or in CTCL^22^ the chromosomal duplication must have occurred at an early stage of T-cell development. Moreover, chromosomal breaking points in CTCL contain RAG heptamer sequences, highlighting the role of RAG in malignant transformation.^23^

In addition to the early precursor hypothesis, malignant transformation of multiple cells in an inflammatory infiltrate could provide an alternative explanation of the observed clonotypic heterogeneity in MF.^24^ *Staphylococcus aureus* often present in skin microbiota, might provide an antigenic drive for MF. This hypothesis was supported by findings of a higher than expected usage of Vβ segments involved in recognition of staphylococcal superantigens (e.g. TRBV20 or TRBV5.1).^12,25,26^ We could not confirm those observations; on the contrary, we found that MF clonotypes including those shared between patients, contain Vα and Vβ segments that are found at a very low frequency in peripheral blood or in the inflamed skin (e.g. pseudogenes TRAV11, TRAV28, TRAV31, TRBV12-1, TRBV22-1).^27,28^ We hypothesize that the putative increased frequency of pathogen recognizing Vβ usage may be due to presence of reactive T-cell in the sample, which was minimized in our material which was microdissected and enriched in neoplastic cells.

We have demonstrated that probe capture WES/WTS is a useful and straightforward approach to the analysis of clonotypic composition in MF. Our data show existence of multiple malignant TCR clones in CTCL. The significance of clonotypic heterogeneity for disease prognosis and clonal evolution during the course of the disease remains to be investigated.

## Materials and Methods

### Sample collection and storage

Ethical approval was obtained from the Health Research Ethics Board of Alberta, Cancer Committee HREBA.CC-16-0820-REN1. After informed consent, 4mm punch skin biopsies were collected from patients and embedded in the optimal cutting temperature (OCT) medium at - 80°C. 10 ml of blood was collected and Ficoll was used to isolate peripheral blood mononuclear cells (PBMC) that were subsequently resuspended in 50% of Dulbecco’s modified eagle medium (DMEM), 40% Fetal bovine serum (FBS) and 10% DMSO and frozen in liquid nitrogen until further use.

### Cryosectioning and laser capture microdissection (LCM)

10 μm sections of the skin biopsies frozen in OCT were collected on 2 μm polyethylene naphthalate (PEN) membrane slides (cat# 11505158, Leica Microsystems, Wetzlar, Germany). The slides were then stained using hematoxylin and eosin stains to identify the tumor cells. The microdissected tumor cell clusters were collected in RLT buffer (cat# 79216) (Qiagen, Hilden, Germany) and used for simultaneous DNA/RNA isolation using AllPrep DNA/RNA micro kit (cat# 80284, Qiagen, Hilden, Germany). Isolated DNA was preamplified using REPLI-g single cell kit (cat# 150343, Qiagen, Hilden, Germany).

### Sample preparation for whole exome sequencing (WES)

1 μg of DNA measured using Qubit™ dsDNA HS assay kit (cat# Q32851) (Thermo Fisher, Massachusetts, United States) was sheared at a peak size of 200 bp using Covaris S220 focused-ultrasonicator (Covaris, Massachusetts, United States). Sheared DNA was end-repaired, adapter ligated and indexed using NEBNext^®^ Ultra™ II DNA library prep kit for illumina (cat# E7645S) (New England Biolabs, Massachusetts, United States). Prepared libraries were hybridized with biotin cross-linked RNA baits (SSELXT Human All exon V6 +UTR) (Agilent Technologies, California, United State) at 65 °C for 24 hours. Few of the samples were also used for hybridization with custom probes designed to target V and J TCRα, TCRβ and TCRγ sequences. These custom probes were combined with the current SSELXT Human All exon V6 +UTR kit (Custom + SSELXT Human All exon V6 +UTR) to improve the overall efficiency of the capture protocol in identifying TCR clonotypes. Hybridized DNA was captured using Dynabeads™ MyOne™ streptavidin T1 (cat# 65601) (Thermo Fisher, Massachusetts, United States). Captured DNA was amplified using KAPA library amplification kit with primers (cat# 07958978001) (Roche Diagnostics, Risch-Rotkreuz, Switzerland). Prepared DNA libraries peak size was verified using 2100 Bioanalyzer, (Agilent Technologies, California, United State). The DNA libraries were sequenced on an Illumina HiSeq 1500 sequencer using paired-end (PE) 150 reads.

### Sample preparation for whole transcriptome sequencing (WTS)

10 ng of total RNA quantified using Qubit™ RNA HS assay kit (Q32852) (Thermo Fisher, Massachusetts, United States) was used for rRNA depletion (E6310) (New England Biolabs, Massachusetts, United States). rRNA depleted samples were used for cDNA synthesis and prepared to be sequenced using NEBNext^®^ Ultra™ II directional RNA library prep kit for illumina (E7760) (New England Biolabs, Massachusetts, United States). Prepared cDNA libraries peak size was verified using 2100 Bioanalyzer, (Agilent Technologies, California, United State). The cDNA libraries were later sequenced on an Illumina HiSeq 1500 sequencer using paired-end 150 reads.

### Data analysis

The sequencing fastq files were analyzed using MiXCR to identify the TCR clonotypes.^29^ For WTS short and long read alignments were included. But for WES data, partial reads were filtered out as these might be the captures of only V or J sequences. The sequencing reads were processed using the GATK4 generic data-preprocessing workflow, then analysed with Titan^30^ to determine copy number aberration and tumor purity using the hg38 Human reference genome. tcR package was used to calculate the inverse Simpson diversity index and identify the overlapping clones.^31^ VJ combination bias was analyzed using VDJtools package.^32^

## Acknowledgement

We would like to thank Dr. Weiwei Wang for his input in the study design. We would also like to acknowledge Mrs. Rachel Doucet and the nursing staff of Edmonton Kaye clinic for their help in sample collection. This study was supported by the grants from the following sources: Canadian Dermatology Foundation (CDF RES0035718), University of Alberta, Bispebjerg Hospital (Copenhagen, Denmark) unrestricted research grant to R.G., and Danish Cancer Society (Kræftens Bekæmpelse R124-A7592 Rp12350).

## Authors contribution

AI designed, performed the experiments, analyzed the data and wrote the manuscript. JP performed CNV analysis and tumor purity calculations. GW provided input with the technical aspects of the experiments and edited the manuscript. TS helped with samples collection and recruitment of patients for the study. RG supervised the experiments, data analysis and edited the manuscript. All authors approved the final version of this paper.

## Supplementary Figures

**Supplementary Figure S1:**
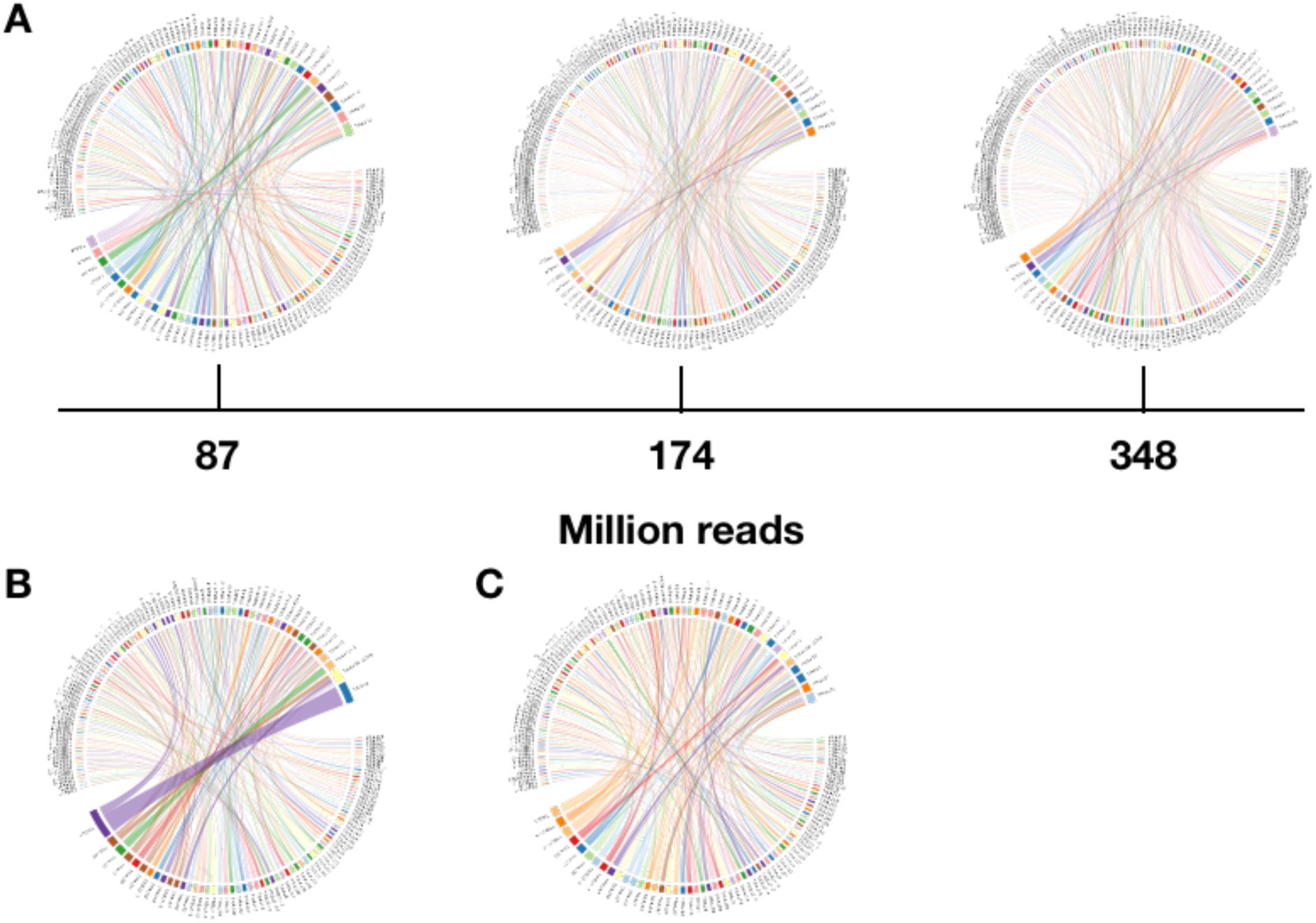
VJ gene usage. A. **A** peripheral blood mononuclear cells (MF5 PBMC) sample was used for whole exome capture and sequenced at a maximum depth of 400 million reads. Sequenced samples were analyzed for V and J gene combination and no preferential combination was identified as oppose to the high frequency VJ combination identified in **(B)** that can be associated with presence of dominant clone in a MF5T sample. **(C)** VJ gene usage of MF5P.

**Supplementary Figure S2:**
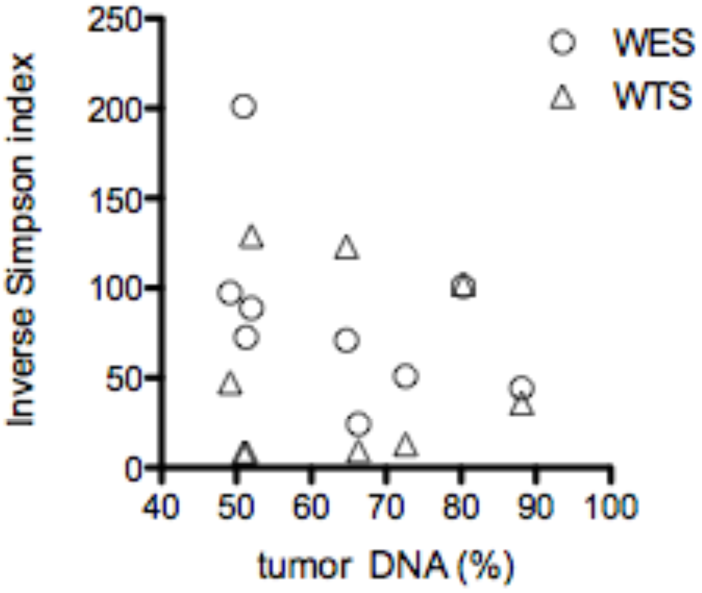
Relationship between inverse Simpson index and tumor enrichment. Microdissected island containing atypical lymphoma cells were subjected to WES/WTS. Inverse Simpson index reflects the TCR repertoire richness. Proportion of tumor-derived DNA in the sample was calculated based on copy number aberration analysis from WES. Symbols represent values for individual samples.

**Supplementary Figure S3.**
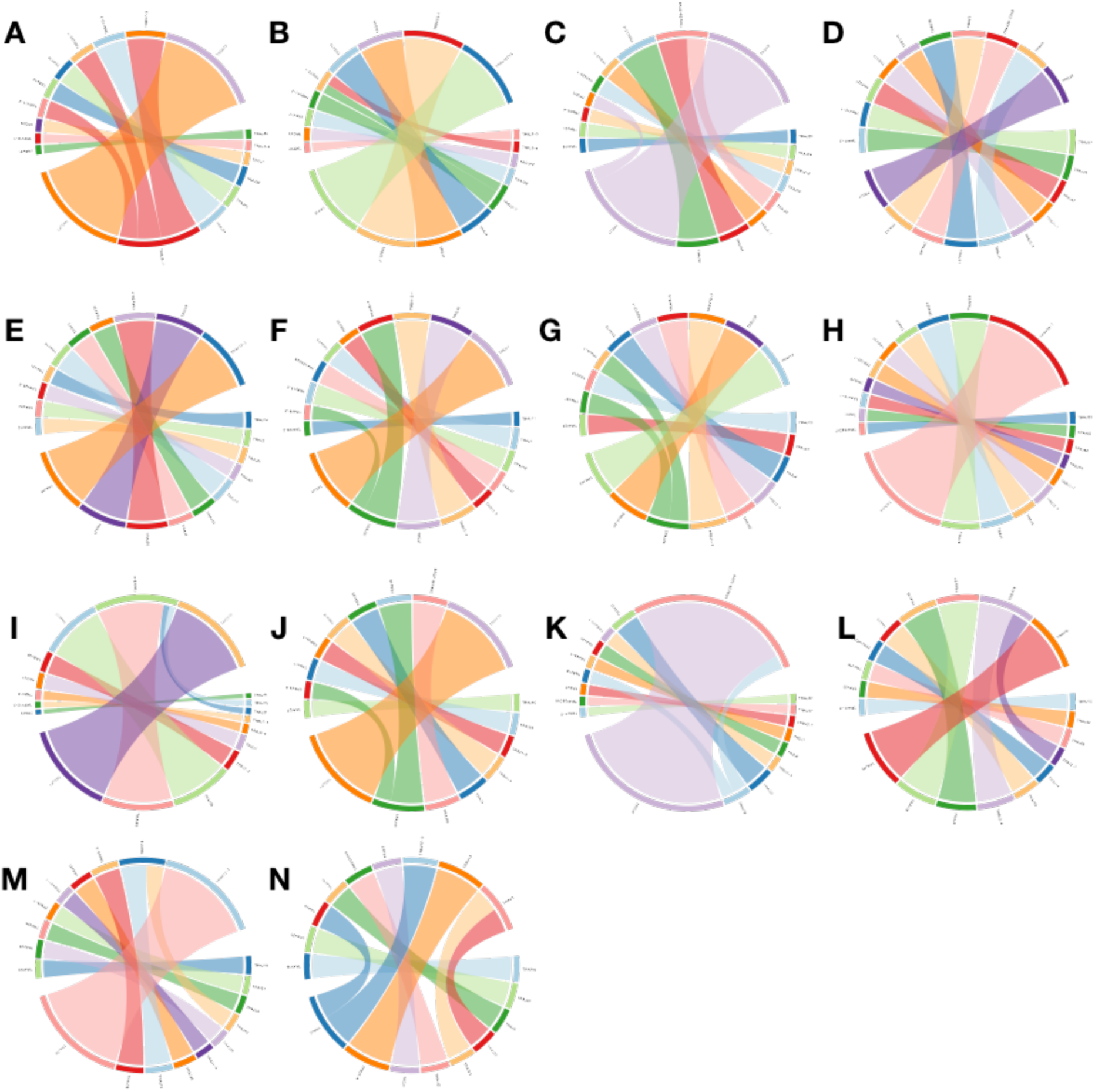
VJ gene usage for top 10 high frequency clonotypes identified using WES in top 10 dominant clones. The samples correspond to those in Fig 3.

**Supplementary Figure S4.**
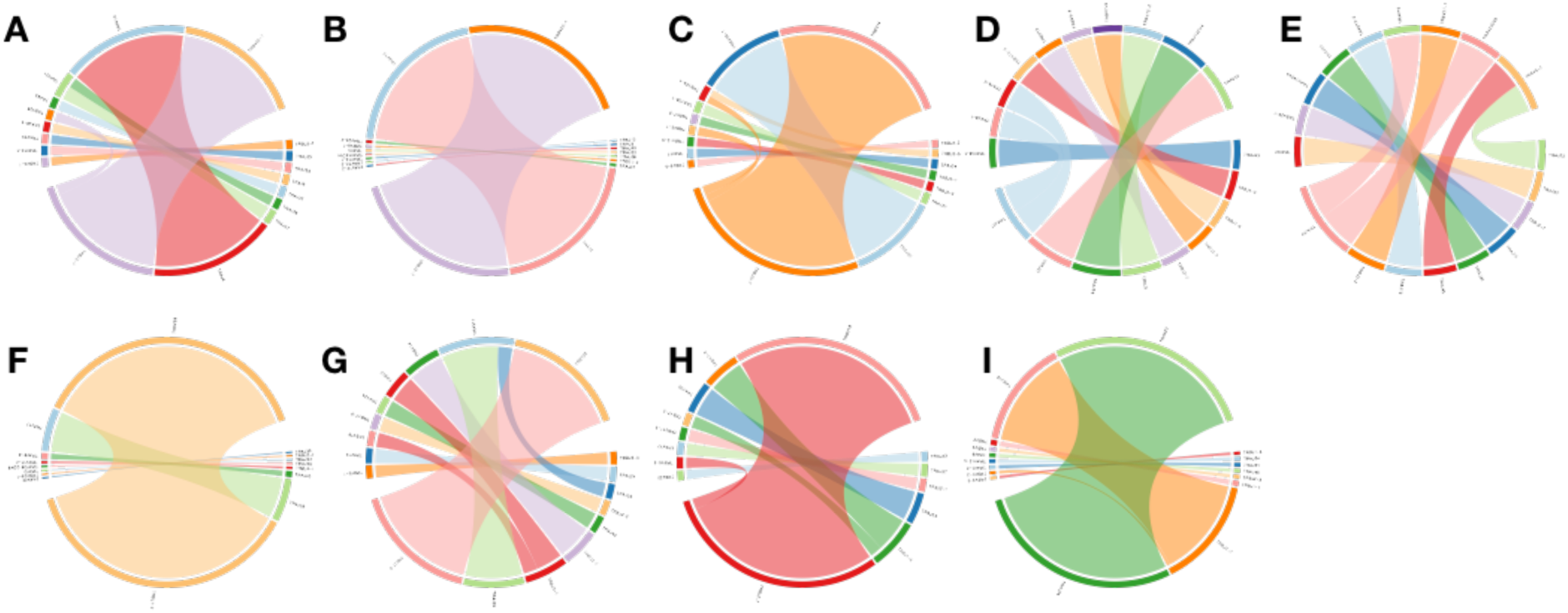
VJ gene usage for top 10 high frequency clonotypes identified using WTS in top 10 dominant clones. The samples correspond to those in Fig 3.

**Supplementary Table S1.** Number of clonotypes identified by WES (DNA) and WTS (RNA) in patients with MF. Samples are annotated as patient number (see Fig 1) with the suffix P (plaque), T (tumor) or PB (peripheral blood mononuclear cells). Normal Lymphocytes are pooled CD4+ cells from 4 healthy donors.

| Samples<br>(DNA) | Number of TCR clonotypes |  |  | Samples<br>(RNA) | Number of TCR clonotypes |  |
| --- | --- | --- | --- | --- | --- | --- |
| | TCRα | TCRβ | TCRγ | | TCRα | TCRβ |
| MF4T | 393 | 95 | 15 | MF4T | 48 | 24 |
| MF4P | 279 | 63 | 7 | MF4P | 86 | 47 |
| MF5T | 160 | 41 | 8 | MF5T | 98 | 148 |
| MF5P | 138 | 39 | 7 | MF7T | 21 | 10 |
| MF7T | 108 | 21 | 4 | MF7P | 29 | 21 |
| MF7P | 109 | 29 | 5 | MF11T | 12 | 4 |
| MF9 | 121 | 22 | 2 | MF11P | 179 | 131 |
| MF10 | 66 | 22 | 2 | MF19T | 6 | 10 |
| MF11T | 157 | 45 | 3 | MF19P | 134 | 84 |
| MF11P | 146 | 45 | 8 | Normal<br>Lymphocytes | 52 | 204 |
| MF17 | 100 | 13 | 3 |  |  |  |
| MF19T | 375 | 81 | 15 |  |  |  |
| MF19P | 141 | 32 | 5 |  |  |  |
| MF23 | 373 | 78 | 11 |  |  |  |
| MF5 PBMC | 130 | 38 | 9 |  |  |  |
| MF11 PBMC | 83 | 15 | 5 |  |  |  |

**Supplementary Table S2.** Percentage of tumor DNA purity and dominant clone frequency for TCR-ɑ, -β or -γ listed. Samples annotated as patient number followed by P for plaque and T for tumor (see figure 2B).

| **Samples** | **Tumor DNA Purity (%)** | **Alpha (%)** | **Beta (%)** | **Gamma (%)** |
| --- | --- | --- | --- | --- |
| MF4T | 72.61 | 5.45 | 19.33 | 74.19 |
| MF4P | 88.13 | 13.68 | 31.25 | 70 |
| MF5T | 64.73 | 4.8 | 10.16 | 64.58 |
| MF5P | 43.11 | 2.28 | 5.76 | 41.66 |
| MF7T | 52 | 5.58 | 8.69 | 66.66 |
| MF7P | 49.18 | 3.59 | 12.5 | 60 |
| MF11T | 66.3 | 15.51 | 17.56 | 84.28 |
| MF11P | 80.35 | 3.9 | 8.82 | 51.72 |
| MF19T | 50.98 | 3.76 | 12.2 | 18.18 |
| MF19P | 51.34 | 10.08 | 9.3 | 20 |

**Supplementary Table S3.**
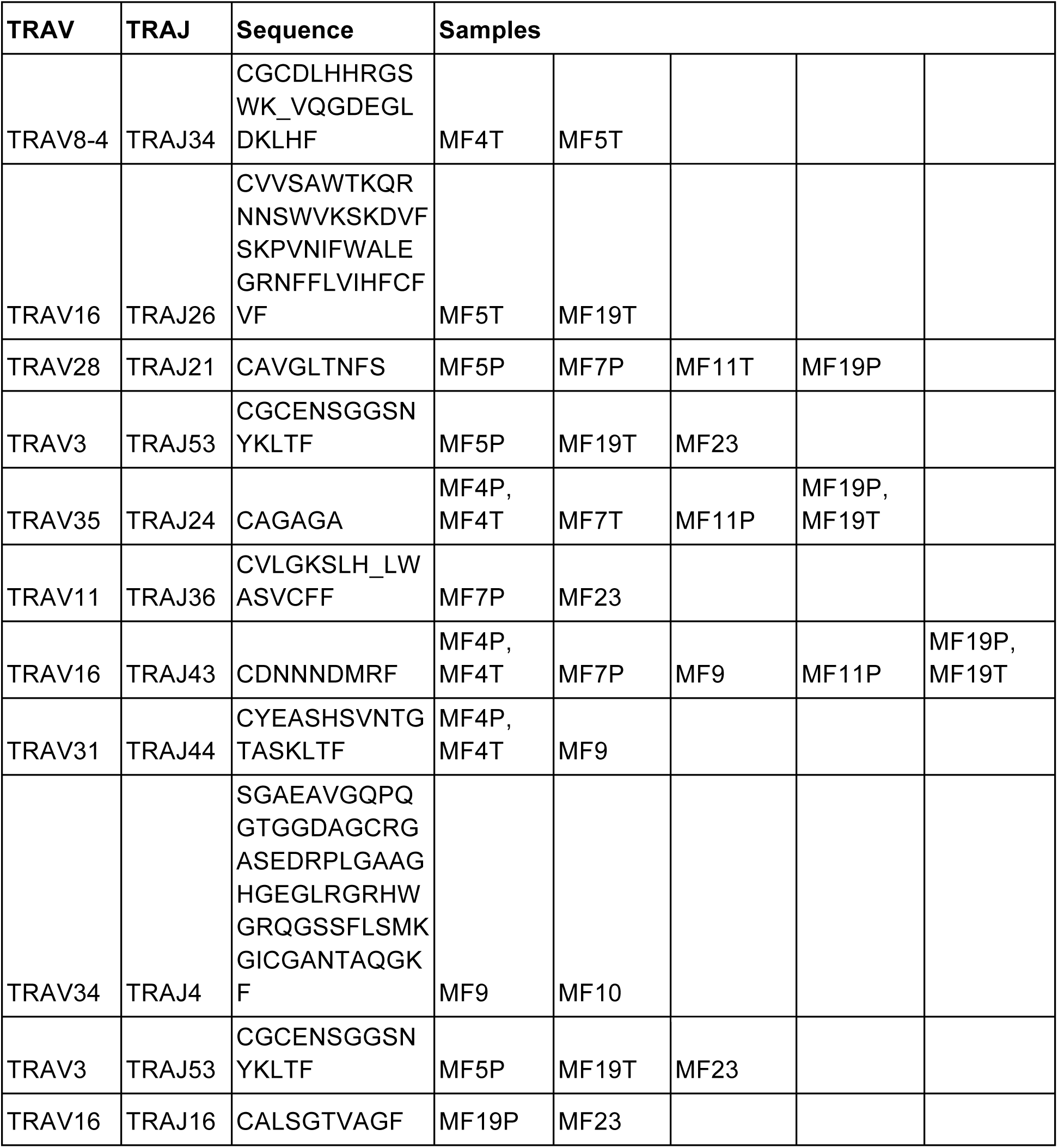
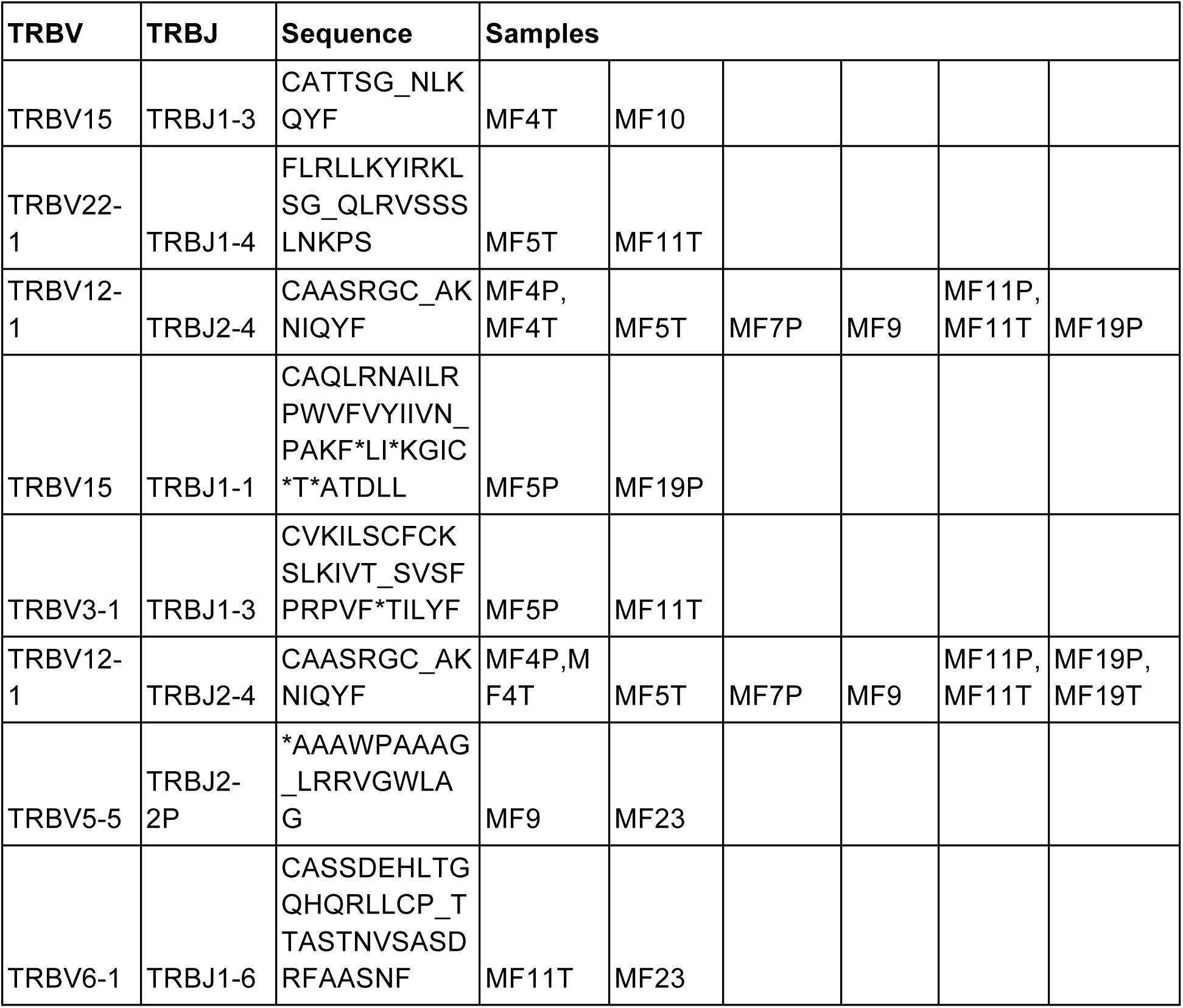
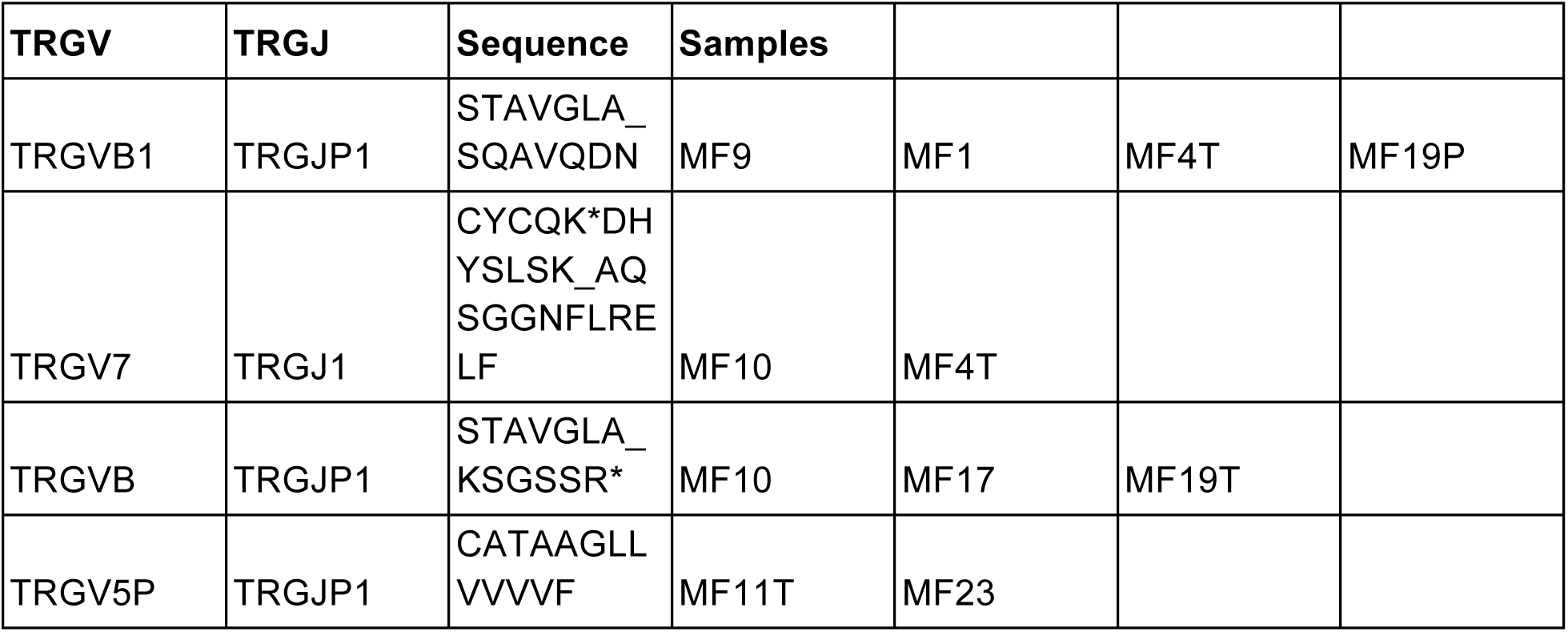
A list of 10 most abundant clonotypes shared across the samples from lesional skin in MF. The samples are annotated as in supplementary table S1.

